# A cost effective high-resolution climbing assay applied to *Drosophila* Parkinson’s and proprioception mutants reveal novel behavioural phenotypes

**DOI:** 10.1101/426544

**Authors:** Aman Aggarwal, Heinrich Reichert, K. VijayRaghavan

## Abstract

Severe locomotor impairment is a common phenotype of neurodegenerative disorders such as Parkinson’s disease (PD). *Drosophila* models of PD, studied from more than a decade, have helped in understanding the interaction between various genetic factors, such as *parkin* and PINK1, in this disease. To characterize locomotor behavioural phenotypes for these genes, fly climbing assays have been widely used. While these simple current assays for locomotor defects in *Drosophila* mutants measure some locomotor phenotypes well, it is possible that detection of subtle changes in behaviour is important to understand the early manifestation of locomotor disorders. We introduce a novel climbing behaviour assay which provides such fine-scale behavioural data and tests this proposition for the *Drosophila* model. We use this inexpensive, fully automated, high resolution assay to quantitatively characterize the parameters of climbing behaviour in three contexts. First, we characterize wild type flies and uncover a hitherto unknown sexual dimorphism in climbing behaviour. Second, we study climbing behaviour of heterozygous mutants of genes implicated in the fly PD model and reveal previously unreported prominent locomotor defects in some of these heterozygous fly lines. Finally, we study locomotor defects in a homozygous proprioceptory mutation (*Trp-γ^1^*) known to affect fine motor control in *Drosophila*. Moreover, we identify aberrant geotactic behaviour in *Trp-γ^1^* mutants, thereby opening up a finer assay for geotaxis and its genetic basis. Our assay is therefore a cost-effective, general tool for measuring locomotor behaviours of wild type and mutant flies in fine detail and can reveal mild motor defects.

**Significance statement:** Fine control of neuronal activity is required for proper motor output. Severe locomotor impairment is a common result of neurodegenerative disorders such as Parkinson’s disease (PD). The fruitfly, *Drosophila*, has been widely used as a model system to study the genetics of these disorders and simple climbing assays have been used to study the behavioural phenotypes of mutations in these genes. Here we introduce a novel, fully automated, high resolution climbing behaviour assay and use this assay to characterize climbing behaviour in wild type flies and in various fly mutant lines related to PD and defects in proprioception. Our assay is a general tool for measuring locomotor behaviours of flies in fine detail and can reveal very mild motor defects.

## Introduction

Movement such as locomotion is the output of the nervous system of the living beings (Sherrington, 1906) (1). Locomotion entails not only properly functional body structures, but also the neuronal mechanisms giving the output and processing information from various sensory modalities. Locomotion has been studied in detail for decades in a multitude of organisms ranging from simple to complex organisms. A fine control of neuronal activity in a well-orchestrated manner has been shown to be necessary for proper locomotor output. The slightest disruption in this regulation can be highly detrimental to the animal’s ability to move its limbs in a coordinated fashion.

In *Drosophila*, negative geotaxis is an integral part of its locomotor behaviour and has been studied for over a century (2). Climbing assays have been used to identify and study molecules involved in fly models of fine motor control (3), Alzheimer’s disease (4), Parkinson’s disease (5), ageing (6) and motor function degenerative disorders such as spinal muscle atrophy (7). Although these studies have unveiled a plethora of valuable information, one major aspect which has been overlooked is that all these results are based on mechanical stimulation of the flies. For example, most common climbing assays employ tapping down of the flies onto hard glass/polystyrene surface of test tubes (6, 8). This regime of coercing flies to the bottom of the test tubes indeed is very effective for multiple repeats but could also induce physical stress and trauma to the flies (9). As it is known that exposure to physical stress can alter the behavioural output of an animal, assays implementing tapping of flies could miss out fine locomotor differences, if not the most prominent ones.

In his classical work, Carpenter studied various combinations of phototaxis and geotaxis in flies (2). He postulated that light induces locomotion whereas gravity induces directionality in freely moving flies. Positive phototaxis with regard to negative geotaxis have been further studied in the RING assay by the Benzer and other labs, with mechanical agitation being an intricate part of these assays (8, 10). Mechanical stimulation could be a deterrent in the accurate measurement of behaviour. Carpenter used mechanical stimulation only when flies showed little to no locomotion and it is known that the flies have bouts of activity interspersed with non-active periods resulting in activity duration of not more than 40% of the total time (11). Thus, there is spontaneous locomotor activity in fruit flies which could be studied without any aggressive stimulation. Hence, a climbing assay which addresses this issue could be useful in tracking even subtle behavioural differences.

Multiple behavioural assays have been developed decades ago to assay negative geotactic behaviour of flies, ranging from the simple flies in a cylinder (2) to counter current RING assay (8) and geotaxis maze (12). However, there has not been any significant advancement in climbing behaviour assays since then. Traditionally, fly climbing assays have been manual and labour intensive. Usually, the climbing behaviour is scored by keeping track of the flies visually and scoring for a minimum distance climbed in a fixed time duration (10). Along with being time consuming, these assays inadvertently introduce inconsistency in the results. A mechanised system with minimal human intervention for measuring the behaviour would be ideal for getting robust and consistent behavioural output. Not only could the measuring apparatus be automated, data analysis could also be computerised, thus making the output even more reliable and alleviating the need of countless hours of manually keeping track and timings of the flies. Here we describe a novel assay for fly climbing behaviour which does not cause mechanical agitation and is automated for behaviour and data analysis. The assay can be used to analyze behaviour from a single fly or multiple flies without making any change in the setup. We use this fully automated, high resolution assay in single fly assay mode to quantitatively characterize climbing behaviour in wild type flies as well as in two mutant contexts: mutants of genes implicated in the fly PD model and a mutant in proprioceptory structures.

The *parkin*, PINK1 and *Lrrk* genes are some of the most rigrously studied genes in fly model of PD. Although, mutations in *parkin* and PINK1 are known be autosomal recessive, heterozygous mutations in these genes are thought to enhance the risk for PD early onset (13). Here, we use *park^25^*/+ flies to test climbing specifically in heterozygous *parkin* mutants and we also test a commonly used experimental control PINK1^RV^ which is a revertant allele for PINK1 mutation (14). Mutations in *Lrrk* cause late-onset autosomal dominant PD and are one of the strongest risk factor in sporadic PD (15). Studies in flies involving *Lrrk* gene have looked at behavioural manfestaions of the PD phenotype in homozygous condition. We examined the behaviour of heterozygous *Lrrk^ex1^*/+ mutants and studied effect of *park^25^* mutations in trans-heterozygous state with *Lrrk^ex1^* mutation. Behavioural pehnotypes in these PD fly models are not reported in young heterozygous flies and using our newely developed assay, we observed severe behavioural phenotypes in these fly models of PD at very young stage in their life cycle.

Further, to test the ability of this assay to reveal subtle changes in climbing abilities of the flies, we studied the climbing locomotor behaviour of a proprioceptory mutuant, Trp-γ^1^. Trp-γ is a TRPC channel which is known to be expressed in mechano-sensory neurons of thoracic bristles and femoral chordotonal organ. The *Trp-γ^1^* mutant has defects in fine motor control, but otherwise no major locomotor defects (3). Here we identify aberrant geotactic behaviour in *Trp-γ^1^* mutants, thereby opening up a finer assay for geotaxis and its genetic basis.

## Materials and methods

### 1) Fly culturing

Flies were grown at 23°C with 12/12 light/dark cycle. One day old flies were collected and both-sex cohorts were maintained on standard corn meal agar vials for the assay. Climbing behaviour was performed on 3-4 day old flies. *Canton-S* flies were used as wild type. CS, w^1118^;;, PINK1^RV^;;, w^1118^;; Trp-γ^1^ and w*;; *Lrrk^ex1^* /TM6b,TB flies were obtained from Bloomington *Drosophila* Stock center. w^−^;; *park^25^*/TM3-Ser flies are used as described already (16). Males from w^-^;; *park^25^*/TM3-Ser and w*;; *Lrrk^ex1^*/TM6b,TB were crossed with virgin females of w^1118^;; to generate w^1118^;; *park^25^*/+ and w*;; *Lrrk^ex1^* /+ flies, respectively. Trans-heterozygote flies of genotype w*;; *Lrrk^ex1^/ park^25^* were obtained by crossing males of w*;; *Lrrk^ex1^* /+ with virgins females from w^−^;; *park^25^*/TM3-Ser.

### 2) Climbing setup

The behaviour cassette (fig 1B) design is modified from a previously published design (17) and was custom built with three sections: top, middle and bottom, which were held together with four screws. The top and middle section were fabricated using transparent acrylic sheet of 3mm thickness. A 6mm diameter circular slot was drilled in the top section for introducing flies into the cassette. The middle section contained a 110mm × 10mm × 3mm (length × breadth × height) slot which was used as climbing arena for the flies. The bottom section was made of translucent white acrylic sheet to act as a diffuser for the background light for uniform illumination.

**Figure 1.**
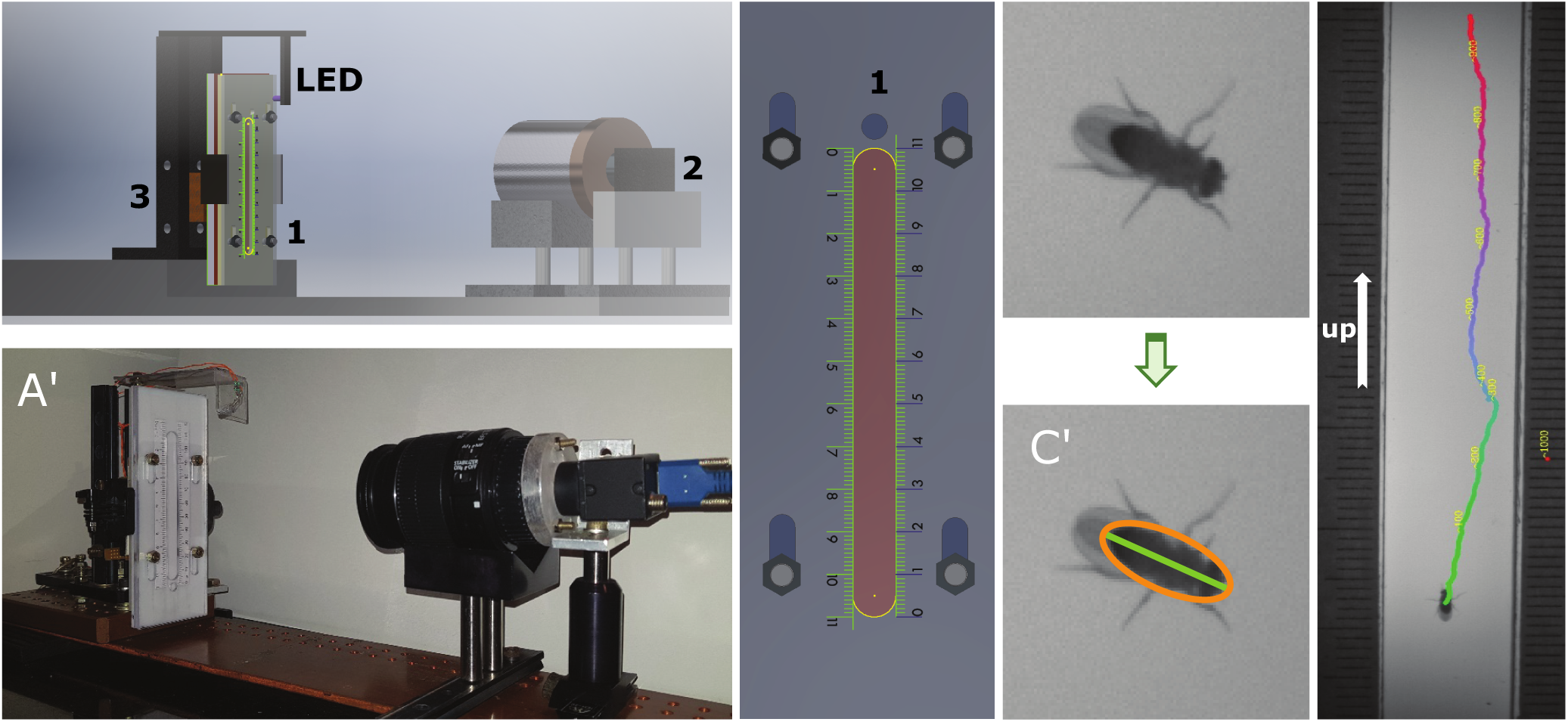
Fly climbing setup. (A) 3D model of the fly climbing/experimental setup consisting of a climbing cassette (1), camera (2), cassette rotating mechanism (3), and a UV LED light. (A’) Actual image of the climbing setup. (B) Schematic of the behaviour cassette in which (1) depicts the hole from where the experimental fly is introduced in the climbing arena. (C) Top view of a wild type, *Canton-S* (CS), male Drosophila, during one of the recorded climbings. (C’) Representative segmentation of the same fly in the same video frame, with the orange oval representing the automatically detected body contour and green line depicting fly body length. (D) Raw frame of a fly climbing up in the arena, superimposed with its track after post-hoc calculation by FlyConTra software. Green to red colormap indicate the position of the fly in the track, with Green and red color indicating the start and end position of the fly, respectively. Numbers in yellow represent the frame number w.r.t position of the fly in the track, detected by FlyConTra, at that particular time point.

Backlight: The behaviour cassette was uniformly illuminated from back using a custom built infrared (IR) light source. The IR light source was built using 8xIR LEDs (Osram SFH4550) connected in series and powered by a standard 15V switch-mode power supply using current controlled circuit. Custom built light guide panel system was used to provide uniform illumination to the climbing arena.

Vertical rotation mechanism: The climbing behaviour setup is based on the vertical rotation of the behaviour cassette. An Arduino controlled servo motor (Futaba S3003, https://www.futabarc.com/servos/analog.html) based mechanism was implemented for vertical rotation (in one plane) of the behaviour cassette. The servo rotates at 180 rpm with a stationary period of 15 seconds per 180° rotation. The rotation speed of the arena is extremely slow as compared to fly’s turning speed (18). The backlight and the behaviour cassette is mounted on the servo motor using a retractable 50mm x 35mm custom made clamp. The retractable clamp provides ability to quickly disengage the cassette from the rotation mechanism for changing the cassette.

### 3) Arena details

A sliding mechanism in the behaviour cassette allows the fly entry slot to be superimposed on the climbing arena. Single fly is gently tapped into the climbing arena from the fly vial without anaesthesia. Further, behaviour cassette is clamped to the servo to allow vertical rotation of the cassette. The assay is started soon after the fly’s entry into the arena.

### 4) Image capturing

Fly behaviour is captured at 250FPS using a Pointgrey camera (13Y3M) with a Canon 18-55ES lens at 55um/pixel resolution. The lens aperture is maximized to increase the light capture. The depth of field of the lens assembly allows capturing good resolution images of a fly climbing on either top or bottom section of the arena. A constant area of the arena, 70mm × 10mm, was imaged which enabling analysis of the climbing behaviour at high spatio-temporal resolution.

### 5) FlyConTra software

a. Data capturing: Imaging is done using custom FlyConTra (**Fly Con**tour based **Tra**cker) program, written in *Python*, which employs motion detection for image capturing. Image capturing is started immediately after the cassette is clamped on to the rotation assembly. Motion detection in FlyConTra initiates image capturing if there is any movement, within the defined thresholds, in the arena being filmed. Algorithms deployed in FlyConTra help in capturing images only when the fly is moving in a stationary arena, thus minimizing the amount of data captured. Image analysis for fly climbing behaviour is done by FlyConTra post-hoc.
b. Image analysis: FlyConTra detects the fly based on its shape and contrast with the background. Parameters defining the body shape and contrast can be adjusted in the software for rough approximation of fly shape. FlyConTra calculates the exact fly shape parameters for each fly by analysing the dataset of that fly. Fly is detected in each frame and the coordinates of its position in sequential frames are saved in a corresponding file. This positional information is then used to calculate the further climbing parameters such as average speed, number of tracks, distance travelled, etc.

### 6) Parameters calculated

We used fly track as a basic unit to measure locomotion of flies in the arena. A track is defined as the path treaded by a fly when it moves continuously for a distance greater than one body length unit (BLU) without stopping for more than one fifth of a second at a time. For example, if a fly, with BLU approximately 2.53mm, moves a distance of 15mm, but stops for 0.3 seconds after moving 9mm, FlyConTra would calculate that as two tracks. The following parameters of fly locomotion were calculated from the data captured from various flies of a given strain:

a. Total number of tracks: Total number of tracks of a genotype in a given time duration is the mean of number of tracks climbed by each fly of that genotype.
b. Total distance travelled: Distance travelled by a fly is equal to the sum of all track lengths in BLUs in a given time duration. For a genotype, total distance travelled in a given time is calculated as the mean of total distance travelled by each fly of that genotype.
c. Track duration: Duration of a track is the median of the duration, in seconds of each track, once the fly initiates movement. For a genotype, the duration of tracks, in a given time, is calculated by taking the mean of median track duration of each fly of that genotype.
d. Average speed: For each track, the speed of the fly is calculated as the average instantaneous speed for that track. Further, mean speed of a fly is given by the mean of speeds for all tracks. Finally, for a genotype, average speed is calculated as the mean of mean speed of each fly.
e. Average track straightness: The track straightness is the coefficient of determination, r2 value, of the linear regression model of the fly track. For a genotype, the average track straightness is calculated as the mean of coefficient of determination of all tracks of a fly.
f. Geotactic Index: Geotactic index is the measure of a fly’s ability to sense and act apropos of gravity. Each track in which the fly moves against gravity (from bottom towards the top of the arena) is scored −1. Similarly, each track where fly moves along gravity (from top to bottom of the arena) is scored +1. Tracks where the fly does not show any vertical displacement above given threshold, are scored zero (0). Geotactic index is calculated as:

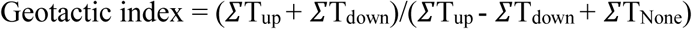 Where:

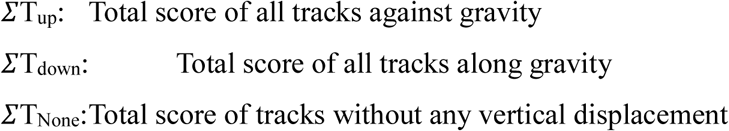

### 7) Statistical analyses

Statistical analysis of all raw data was done using GraphPad Prism (San Diego, USA). Graphs were plotted using matplotlib library in *Python*. For comparison between two genotypes, unpaired t-test (for normal distribution) or Mann-Whitney test (for non-normal distribution) was used. For comparison between more than two genotypes, one-way ANOVA followed by Tukey’s *post-hoc* test (for normal distribution) or Kruskal-Wallis test followed by Dunn’s *post-hoc* test (for non-normal distribution) was used. For time series data, two way repeated measures ANOVA followed by Tukey’s *post-hoc* test was used. Numerical data is reported as mean ± SEM.

## Results

### Automated analysis of locomotor climbing behaviour in *Drosophila*

Advances in image capturing and analysis techniques during the last couple of decades have made it possible to obtain detailed quantative insights into many aspects of fly locomotor behaviour. Using digital, high speed image capturing technology and analysis we developed a new automated assay which gives us information of fly climbing behaviour with high resolution (fig 1A, 1A’). The setup consists of a fly behaviour cassette (fig 1B) mounted on an automatically rotated cassette holder. The arena, big enough for a fly to move freely (110mm × 10 mm × 3 mm, fig 1B), was carved inside the behaviour cassette and mounted on an automated servo motor. The servo (fig 1A), controlled electronically, gives flies 15 seconds per rotation to climb in the arena. The rotation speed of the arena was not more than 60°per second, much less than a fly’s turning speed (18). The arena was uniformly backlit via infrared light, enabling imaging of freely moving flies in a well illuminated space. An ultra Violet LED was placed towards the top of the cassette (fig 1A). Single fly behaviour was recorded for at least 5 minutes for multiple rotations per fly via imaging and analysed by tracking the fly in the images (fig 1C). The fly was detected on basis of its body’s elliptical shape along with contrast from the background using custom developed FlyConTra software (fig 1C’). Fly track, the basic unit to define the locomotor behaviour of the flies, is defined as the path treaded by the fly when it moves more than one body length continuously without stopping for more than 1 second at a time (fig 1D, supplementary movie 1a). Fly locomotor parameters were calculated from the tracks recorded. Fly behaviour parameters were analysed on a per minute basis as well as for a total of 5 minutes to access the fly activity bouts which also depend upon the time spent in a new arena (11). Time series analysis of locomotion parameters enable us to compare fly behaviour as the fly starts to acclimatize in the arena.

### Locomotion parameters of climbing wild type flies

In order to validate our automated image capturing and analysis setup of a fly in the climbing arena and establish locomotor baseline parameters for further studies with mutants, we first determined the locomotion parameters of climbing wild type flies. In this validation study the following climbing parameters were characterized: number of tracks, average track duration, total distance travelled, average speed and path straightness.

#### Number of tracks

The number of tracks climbed by flies is the direct measure of locomotor behaviour. To estimate the motivation to climb, the total number of tracks were quantified for each fly. Climbing for 38 CS flies was measured. The behaviour cassette rotates at 3.33RPM, a total of around 17 times in 5 minutes (Fig 2A). CS flies climbed 3.42±0.16 tracks (mean±SEM) in the 1^st^ minute and slowed down to 2.92±0.12 tracks per minute for rest of the time (fig 2A’, Supplementary data 2). The decrease in locomotion could be attributed to fly’s acclimatization to the arena. Over all, wild type flies walked for 15.58±0.36 tracks in the first 5 minutes after they were introduced in the climbing arena (fig 2A). The average number of tracks climbed by the wild type flies is comparable to the number of time the behaviour cassette rotates. This result shows that CS flies tend to climb for one track with every rotation of the cassette.

**Figure 2.**
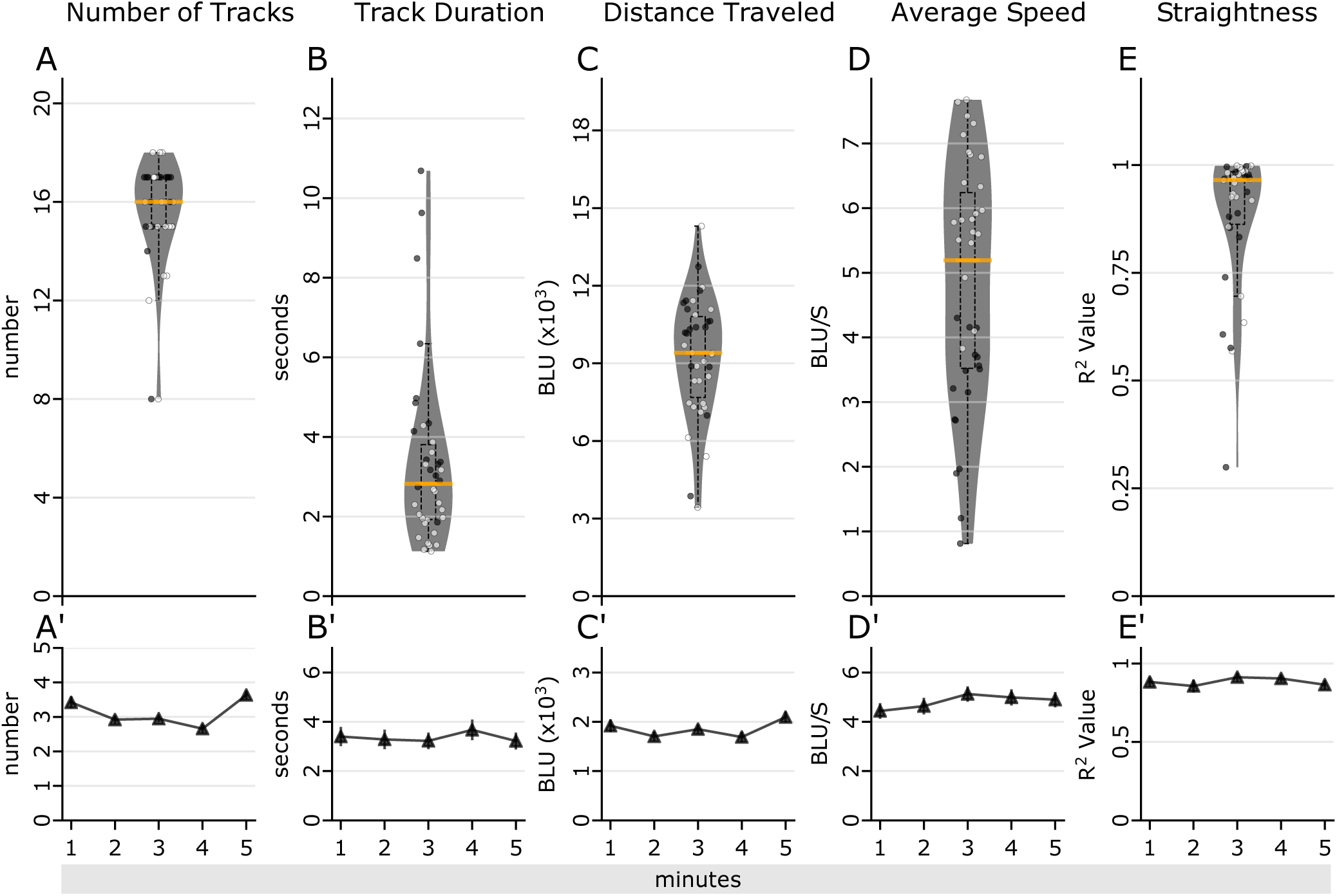
Quantification of general climbing parameters of CS flies. (A) Average number of tracks, (B) Average duration of track, (C) Total distance, (D) Average speed of the fly in climbing and (E) Average path straightness for the paths climbed by the fly, represented by open and closed circles for males and females, respectively. (A’-E’) Time series analysis of each parameter in (A-E) respectively. CS flies on an average climbed 15.58±0.37 tracks (A) with mean track duration of 3.33±0.36 seconds (B) and covered total distance of 9259.30±370.03 BLU (C) with an average speed of 4.81±0.30 BLU/s (D). The average path straightness was 0.89±0.03 (E). The values on per minute analysis did not change much with increase in time spent in the assay (Supplementary data 1, 2). (CS: n>15, for males and females each). The gray area in (A-E) represent the range of full data sets, with median represented as the orange line. Width of the plot indicates of the density of data points. Box inside the colored area correspond to the data within *q1* and *q3*; the extended dotted line represents the data within (*q1*-1.5×IQR): (*q3*+1.5×IQR) range, where *q1* and *q3* are the first and third quartile, respectively; IQR:=*q3−q1* is the interquartile range. BLU: body length unit.

#### Duration of track

Once a fly initiated climbing, the duration of the track was measured. This is a measure of fly’s ability to sustain walking. In the first minute, mean track duration for wild type flies was 3.40±0.38 seconds, which did not vary significantly even after the fly spent more time in the arena (fig 2B’, Supplementary data 2). Overall, wild type flies spent 3.33±0.36 seconds per track (fig 2B, Supplementary data 1). The mean track duration for wild type flies never exceeded 10 seconds, suggesting that 15 seconds is sufficient for a fly to climb in the arena.

#### Total distance travelled

Next we determined the overall locomotor ability of a fly, by calculating total distance travelled by the fly. Using body length unit (BLU) as the basic unit for distance measurements, total distance travelled for each fly was normalized to the body length. In first minute of introduction of the fly into the arena, CS flies travelled 1919.73±122.63 BLUs, which did not vary much with the amount of time spent by the fly in the arena (fig 2C’, Supplementary data 2). In 5 minutes, on an average a fly walked a total distance of 9259.30±370.02 BLUs (fig 2C, Supplementary data 1).

#### Average speed

Instantaneous speed of the fly was determined by measuring the distance travelled by the fly between two consecutive frames. Subsequently, mean speed of a fly in one track was taken as the mean of instantaneous speeds of that track. For one fly, average speed was calculated by taking the mean speed from the speed of each track of that fly. Further, for a given genotype, the average speed was calculated by taking mean of the mean speed from each fly. For the first minute, CS flies moved with a speed of 4.44±0.31 BLUs per second and moved with the similar speeds beyond first minute into the arena (fig 2D’, Supplementary data 2). Overall, the mean speed of CS flies over 5 minutes was found to be 4.81±0.30 BLUs/s (fig 2D, Supplementary data 1).

#### Track straightness

Finally, we calculated the straightness of the path traversed by the fly which is a measure of gross motor control and body orientation. Track straightness is calculated by drawing a regression model of the track and calculating the coefficient of determination, r^2^, for that regression model. The value of r^2^ indicates the track straightness, higher r^2^ value correlates to a straighter track. Average track straightness of a fly is calculated by taking the mean r^2^ for all paths traversed by that fly. For wild type flies, average track straightness for first minute into the assay was 0.88±0.029 and was similar for tracks climbed by the fly in the later minutes (fig 2E’, Supplementary data 2). The average track straightness for all the paths climbed by the fly in 5 minutes was 0.89±0.03 (fig 2E, Supplementary data 1).

#### Sexually dimorphic climbing behaviour parameters

Locomotor differences within the population, based on the sex of the flies, were also characterized for the climbing parameters (males and females represented as open and closed circles, respectively (see fig 2). Although, the number of tracks for wild type females were not significantly different from wilt type males (15.85±0.57, 15.98±0.41, p=0.47, Mann-Whitney test, fig 2A), the average track duration of females was significantly higher than males (5.00±0.32, 3.21±0.27, p<0.0001, Mann-Whitney test, fig 2B). The total distance covered by female wild type flies was similar to males (9749.19±520.50BLUs, 9494.04±315.98BLUs, p=0.67, unpaired t-test, fig 2C, Supplementary data 1), but the average speed of females was significantly lower than males (2.63±0.14, 5.02±0.26, p<0.0001, unpaired t-test, fig 2D). Also, the path straightness of females was slightly lower than males (0.88±0.02, 0.93±0.01, p=0.034, Mann-Whitney test, fig 2E, Supplementary data 1). Finally, per minute analysis reveal that females consistently have higher track duration and lower average speed as compared to wild type males (Supplementary data 2). These data suggest that inherently, climbing female wild type flies walk slower than male wild type flies, but are similar for various other climbing parameters characterized.

### Locomotion parameters of climbing w^1118^ flies (white eyed flies)

Next, we compared the locomotion parameters of climbing wild type CS flies to those of w^1118^ flies. White eyed, w^1118^, flies represent one of the most common genetic backgrounds for mutants and transgenic studies. Moreover, since they provide the genetic background for the mutants characterized below, they serve as controls for these mutants.

These control, w^1118^ flies showed a similar number of tracks as compared to the CS flies in total 5 minutes (14.65±0.78, 15.58±0.36, p=0.99, Mann-Whitney test, fig S1A) and time series analysis also did not show any significant differences (Supplementary data 2, fig S1A’). The average track duration of w^1118^ flies was found to be significantly higher as compared to CS flies in total 5 minutes (4.90±0.23s, 3.33±0.36s, p<0.0001, Mann-Whitney test, fig S1B), although per minute analysis revealed the difference in track duration to be different only in first two minutes (Supplementary data 2, fig S1B’). Both, w^1118^ and CS flies travelled similar amount of distance in total 5 minutes (9803.23±599.14BLUs, 9259.30±370.03BLUs, p=0.68, unpaired t-test, fig S1C, Supplementary data 1) and per minute behaviour (Supplementary data 2, fig S1C’). Average speed of w^1118^ flies, as compared to CS flies was less in over all 5 minutes behaviour (3.15±0.14BLUs/s, 4.82±0.30BLUs/s, p<0.0001, unpaired t-test, fig S1D), as well as in per minute behaviour (Supplementary data 2, fig S1D’). Finally, the track straightness of w^1118^, as compared to the CS flies, was low (0.86±0.02, 0.89± 0.03, p=0.0067, Mann-Whitney test, fig S1E) and per minute analysis indicate that the track straightness in w^1118^ flies decreased after first 2 minutes into the assay (Supplementary data 2, fig S1E’).

Thus, while the climbing behaviour of w^1118^ flies is similar to CS wild type flies in some respects such as number of tracks and total distance travelled, they also manifest significant differences in others characteristics *viz*. track duration, average speed and path straightness.

### Locomotor parameters of heterozygous mutants in genes implicated in the fly PD model

Although, current assays for locomotor behaviour in *Drosophila* measure climbing parameters reasonably well (8, 10), it is possible that detection of more subtle changes in climbing behaviour might be important to understand the early manifestation of locomotor disorders. To investigate this, we next studied the parameters of climbing behaviour for several heterozygous mutants in genes implicated in the fly PD model. Climbing behaviour parameters of control w^1118^ flies were compared to those of PINK1^RV^, *park*^25^/+, *Lrrk^ex1^*/+ and *Lrrk^ex1^/park^25^* flies for behavioural differences.

First, we compared the PINK1^RV^ and *park^25^/+* flies with the controls. Total number of tracks climbed by both PINK1^RV^ and *park^25^/+* flies were comparable to the controls in total 5 minutes (15.07±0.92, 13.79±0.90, 16.37±0.93, p>0.99, p>0.99, Kruskal Wallis test, fig 3A) and for per minute data (Supplementary data 2, fig 3A’). Track duration for PINK1^RV^ and *park^25^/+* flies was also comparable to controls for total 5 minutes (4.46±0.18s, 4.07±0.20s, 4.02±0.16s, p>0.99, p>0.99, Kruskal Wallis test, fig 3B) and for per minute data (Supplementary data 2, fig 3B’). Total distance travelled by PINK1^RV^ and *park^25^/+* flies was also similar to controls in 5 minutes (8990.60±680.44BLUs, 7909.31±614.20BLUs, 10632.57±816.34, p=0.71, p=0.37, one-way ANOVA, fig 3C) and for time series (Supplementary data 2, fig 3C’). Average speed of PINK1^RV^ and *park^25^*/+ flies was again comparable to the controls for total 5 minutes (3.13±0.18BLUs/s, 3.28±0.20BLUs/s, 3.64±0.22BLUs/s, Kruskal Wallis test, fig 3D) and for per minute data (Supplementary data 2, fig 3D’). Both PINK1^RV^ and *park^25^/+* did not deviate much from their track, showing track straightness similar to that of the controls for total 5 minutes’ data (0.77±0.03, 0.86±0.02, 0.77±0.02, p=0.12, >0.99 Kruskal Wallis, fig 3E) and for per minute data (Supplementary data 2, fig 3E’).

**Figure 3.**
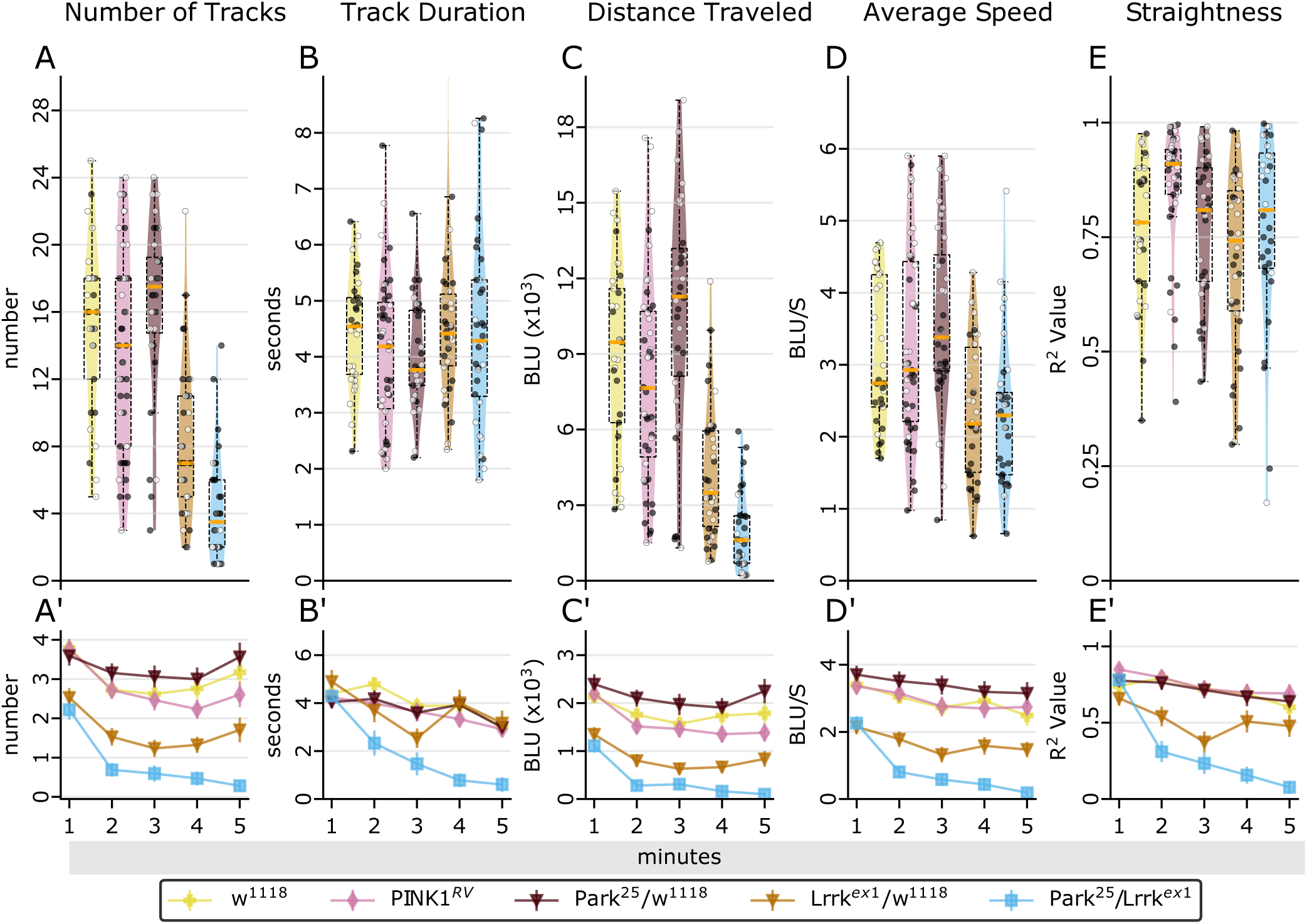
Quantification of various climbing Parameters of Parkinson’s mutants. In total 5 minutes, as compared to controls, *Lrrk^ex1^/+* and *Lrrk^exl^/park^25^* flies climb (A) less number of paths (p<0.0007, p<0.0001, Kruskal-Wallis test) and (C) less distance (p<0.0001, p<0.0001, Kruskal-Wallis test, Supplementary data 1). Average climbing speed of *Lrrk^ex1^ /park^25^* flies is significantly less than controls (one-way ANOVA). Per minute analysis reveal that, w.r.t controls, *Lrrk^ex1^* /+ and *Lrrk^ex1^ /park^25^*show a decreasing trend for all parameters with significantly low values post 1 minute into the assay (Supplementary data 2). Climbing parameters of PINK1^RV^ and Park^25^/+ flies always remain similar to the controls (Supplementary data 2). (w^1118^: n>12, *Lrrk^ex1^* /+: n>15, *Lrrk^ex1^/park^25^:* n>12, Park^25^/+: n>15, PINK1^RV^: n>20, for males and females each). Open and closed circles represent males and females, respectively. (A’-E’) Time series analysis of each parameter in (A-E) respectively. Figure representation similar to figure 2.

Next, we checked the climbing ability *Lrrk^ex1^*/+ and *Lrrk^ex1^/park^25^* flies. As compared to controls, both *Lrrk^ex1^*/+ and *Lrrk^ex1^/park^25^* flies climbed significantly less number of tracks in total 5 minutes (8.32±0.77, 4.25±0.57, p=0.0007, p<0.0001, Kruskal Wallis test, fig 3A) and per minute analysis indicated that both genotypes show a decrease in number of tracks climbed throughout the duration of the assay (Supplementary data 2, fig 3A’). Track duration for *Lrrk^ex1^*/+ and *Lrrk^ex1^ /park^25^* flies was comparable to the controls in total 5 minutes (4.62±0.28s, 4.45±0.30s, p>0.99, p>0.99, Kruskal Wallis test, fig 3B), however per minute analysis showed that track duration of *Lrrk^ex1^/park^25^* flies decrease soon after 1 minute into the assay (Supplementary data 2, fig 3B’). The total distance travelled by both genotypes in 5 minutes was significantly less than the controls (4285.22±452.94BLUs, 1968.40±0.266.30BLUs, p<0.0001, p<0.0001, one-way ANOVAfig 3C) and similarly, per minute analysis showed flies from both genotypes climbing less distance than controls at every minute (Supplementary data 2, fig 3C’). Though, average speed of *Lrrk^ex1^*/+ flies did not show any significant deviation from controls in total 5 minutes, *Lrrk^ex1^*/*park^25^* flies climbed more slowly (2.35±0.16BLUs/s, 2.32±0.18BLUs/s, one-way ANOVA, fig 3D). However, per minute analysis showed that it dropped down significantly after 1 minute into the assay for both, *Lrrk^ex1^*/+ and *Lrrk^ex1^/park^25^* flies (Supplementary data 2, fig 3D’). Finally, track straightness of flies from both genotypes was similar to that of controls in 5 minutes (0.70±0.03, 0.77±0.04, p>0.99, p>0.0856 Kruskal Wallis test, fig 3E) and per minute analysis revealed that the track straightness of flies from both genotypes flies was significantly after first minute into the assay, though *Lrrk^ex1^*/+ flies show straighter tracks during last two minutes of the assay (Supplementary data 2, fig3D’).

Taken together, these findings indicate that, relative to controls, flies from genotypes having mutation in *Lrrk* gene, i.e. *Lrrk^ex1^*/+ and *Lrrk^ex1^/park^25^*, start showing severe locomotor deficits within a minute of introduction into the arena. The other two genotypes, *park^25^/+* and PINK1^RV^ flies do not show any significant departure from controls. The locomotor defects in *Lrrk^ex1^*/+ seem to be excacerbated in the transheterozygote condition with *park^25^* mutation. Although an understanding of the detailed interaction between the two requires further investigation.

### Locomotor parameters in a proprioceptive mutant

Finally, we investigated climbing locomotor defects that result from a homozygous proprioceptory mutation (*Trp-γ^1^*) known to exclusively affect fine motor control in *Drosophila* (3). Previous studies indicate that *Trp-γ^1^* flies have fine motor control defects in locomotion but not any gross motor defect. Thus, overall locomotion in *Trp-γ^1^* flies is not affected much, but large gap crossing is highly impaired (3). We hypothesized that fine motor coordination might be important for climbing. Hence, to further understand the role of proprioceptive neurons expressing *Trp-γ* in motor coordination, we studied climbing of *Trp-γ^1^* flies in greater depth.

Analysis of climbing behaviour of *Trp-γ^1^* flies for a total of the first 5 minutes into the assay showed that, as compared to controls, these flies climbed more tracks (19.85±1.110, 14.92±0.67, p<0.0001, Mann-Whitney test, fig 4A) with a lower average duration of tracks (3.56±0.34s, 4.80±0.20s, p<0.0001, Mann-Whiney test, fig 4B) and travelled a longer distance (11896.05±705.03BLUs, 9627±539.1BLUs, p=0.0117, unpaired t-test, fig 4C) with a greater average speed (4.07±0.23BLUs/s, 3.17±0.14BLUs/s, p=0.0037, unpaired t-test, fig 4D). Track straightness of *Trp-γ^1^* flies was higher than the controls in total 5 minutes (0.90±0.01, 0.81±0.02, p=0.0071, Mann Whitney test, fig 4E). Time series analysis also revealed that, compared to controls, *Trp-γ^1^* flies consistently climbed more number of tracks. For rest of the parameters, per minute analysis showed trends similar to that seen in 5 minutes’ analysis (Supplementary data 2, fig 4A’-E’).

**Figure 4.**
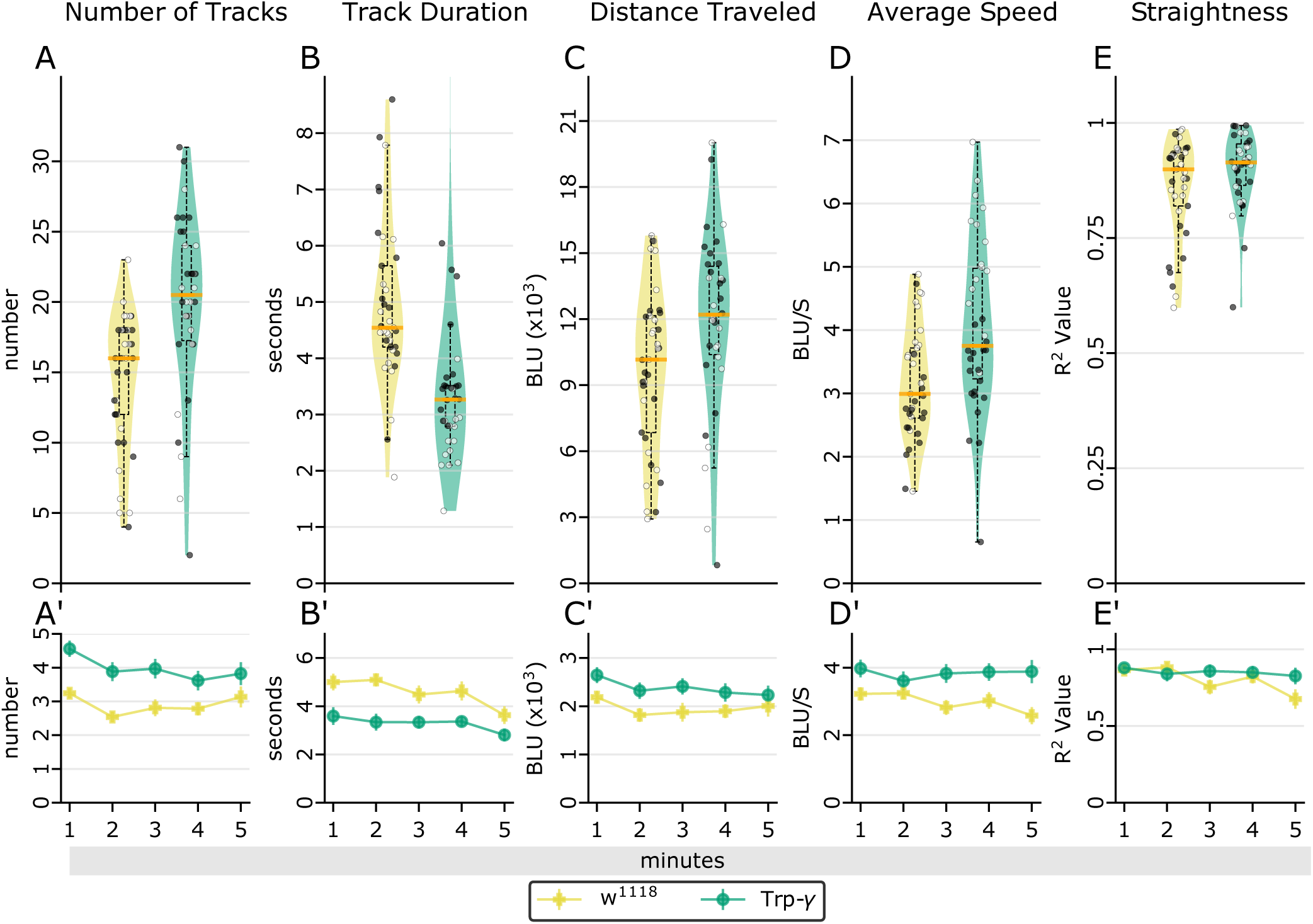
Quantification of various climbing Parameters of Trp-γ mutants. In total 5 minutes, as compared to controls, Trp-γ flies climb (A) more number of paths (p<0.0001, Mann-Whitney test), (B) for shorter duration (p<0.0001, Mann-Whitney test), (C) more distance (p=0.0117, unpaired t-test) and (D) faster (p=0.0016, Mann-Whitney test, Supplementary data 1). Per minute analysis reveal that, w.r.t controls, Trp-γ flies consistently show (A’) more number of tracks, but are otherwise similar to controls (Supplementary data 2). Genotype: Trp-γ: w^−^;; Trp-γ^1^. (w^1118^: n>18, Trp-γ: n>15, for males and females each). Open and closed circles represent males and females, respectively. (A’-E’) Time series analysis of each parameter in (A-E) respectively. Figure representation similar to figure 2.

These results indicate that *Trp-γ^1^* flies have higher motility and higher speeds as compared to the controls while climbing. As *Trp-γ* is expressed in femoral chordotonal organs, it is possible that during climbing precise leg position control is impaired. A detailed gait analysis during climbing in *Trp-γ^1^* flies would be important to understand the role of *Trp-γ* in climbing further.

### Geotactic Index: a measure of graviception in flies

Flies have an innate tendency to climb against gravity. To measure a fly’s response to gravity in our climbing assay, we introduce “geotactic index”(GTI), a measure of fly’s ability to sense and respond to gravity during locomotion. GTI is defined as the sum total of score of all tracks scored for their direction of motion apropos of gravity divided by the total number of tracks climbed by the fly (fig 5B). A track is scored −1 if it shows an ascending climb (Tup), scored +1 if the track shows a descending climb (TDown) and zero if it does not show any displacement of more than 1BLU in vertical direction (Tzero, fig 5A). Thus, a negative or positive GTI shows that the fly is negatively or positively geotactic, respectively.

**Figure 5.**
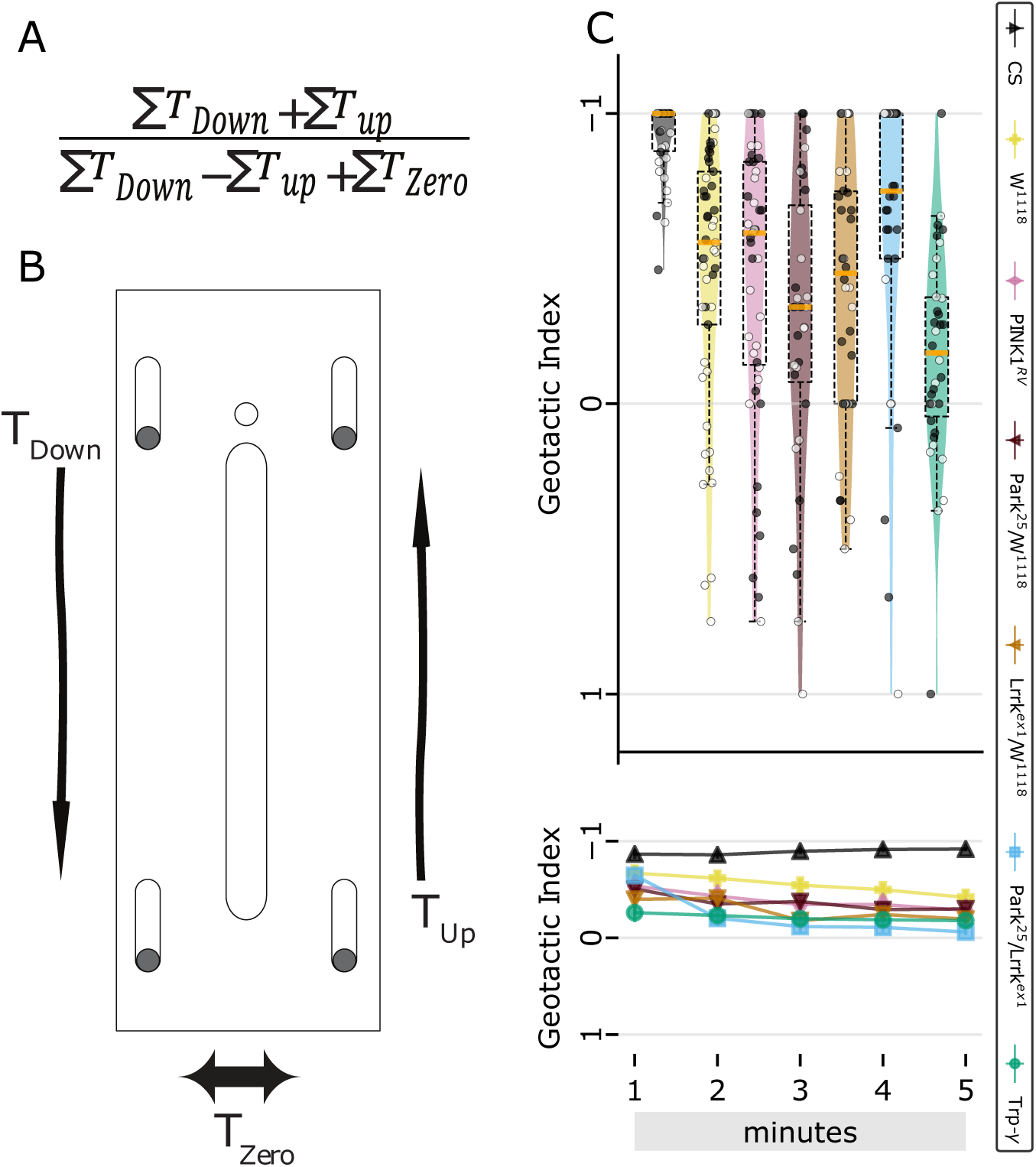
Geotactic Index. (A) Function for Geotactic Index calculation (B) Schematic. (C)The geotactic index of all transgenic flies is lower than CS (p<0.0001 for all genotypes, Kruskal-Wallis test) in the total 5 minutes (Supplementary data 1). (C’) Per minute calculations show geotactic index of *Lrrk^ex1^* /+, *Lrrk^ex1^ /park^25^* and Trp-γ flies consistently lower than CS and control flies (Supplementary data 2). (w^1118^: n>18, Trp-γ: n>15, for males and females each). Figure representation similar to figure 2.

Wild type flies showed a highly negative (−0.92±0.02, fig 5C) GTI in total 5 minutes into the assay. Also, per minute analysis showed that wild type flies do not vary for their GTI even with respect to time (Supplementary data 1, fig 5C’). The control, w^1118^, flies showed less negative GTI as compared to the CS flies in total 5 minutes (−0.45±0.07, p<0.0001, Mann Whitney test, fig 5C) and for per minute data, the GTI increased after 2 minutes into the assay (Supplementary data 1, fig 5C’). *Trp-γ^1^* flies showed a significant increase in GTI, as compared to controls (−0.16±0.06, −0.45±0.07, p<0.0001, Mann-Whitney test, fig 5C) for total 5 minutes and also on per minute basis (Supplementary data 2, fig 5C’).

Overall, these data indicate that a differential response of flies to gravity can be quantified by our geotactic index. Moreover, the results suggest that *Trp-γ^1^* could be involved in the graviception response in flies.

## Discussion

Climbing behaviour in flies has been extensively used to study gross locomotor defects. Widely used climbing behaviour assays employ mechanical startle as a way to induce locomotion in flies. Although highly effective, physical agitation of flies is an aggressive way to induce locomotion in flies. Physical agitation is liable to incorporate undesirable behavioural phenotypes in a fly’s innate climbing ability and mask subtle phenotypes. Also, automating the data collection and analysis could help mitigate the inconsistencies of this traditionally manual, labour intensive task. Therefore, we developed a novel assay to assess climbing ability of flies which hinges upon the fly’s innate response to gravity, *i.e*. negative geotaxis, during climbing. An automated behavioural setup along with robust image analysis is particularly useful in high throughput of this assay.

Advances in computational power have provided huge thrust to computer vision based behaviour analysis methods. Tools ranging from a method, very similar to the one presented here, developed almost a decade ago (19), to more recent machine learning based behaviour annotation tools are being developed extensively (20, 21). Some of these tools, specific for fly limb tracking have greatly contributed to our understanding of fly locomotion at the level of individual leg control (22–25). However, technical challenges limit our ability to quickly screen through large numbers and over long distances using these methods. Our assay mitigates these issues and complements the limb tracking methods already available in the field and provides a way to understand various important factors such as fatigue and geotactic index. Current techniques to analyse climbing are not sensitive enough to pickup these subtle parameters, which could provide crucial insights into the neural mechanisms behind manifestations. Using our method in multiple fly assay mode, one can quickly screen through large number of flies for gross locomotor defects and then can benefit from the already available fine scale gait analysis tools to further narrow down to the potential hits.

This assay could describe some aspects of the fly’s climbing ability in a manner similar to previously used methods, it is important to note that we could gain a lot more information over previously used methods (fig 2). In this study, we describe novel parameters to characterize details of a fly’s climbing behaviour (fig 2A-E), along with a detailed temporal analysis of the same parameters (fig 2A’-E’). Our fine temporal analysis enables an understanding of fly climbing dynamics in accordance with the time spent in the arena, and flexibility in image and data analysis software allows further extraction of various previously undescribed, but important, parameters from the climbing data.

In addition to studying wild type flies, we also characterized climbing behaviour in various fly mutant lines related to PD, *viz. park^25^* and *Lrrk^ex1^*, as well as a commonly used experimental control, PINK1^RV^, which is a revertant allele for PINK1(14). Both the *parkin* and PINK1 genes are implicated in the fly model of PD and although, mutations in *parkin* and PINK1 are known be autosomal recessive, heterozygous mutations in these genes are thought to enhance the risk for PD early onset (13). In addition to using the *park^25^/+* and *Lrrk^ex1^*/+ flies to test climbing specifically in heterozygous state, we also studied *park^25^/Lrrk^ex1^* trans-heterozygous flies. Presence of *park^25^* mutation in conjunction with *Lrrk^ex1^* mutation exacerbates the locomotor defects in fly PD model. Using this trans-heterozygous mutant, we were able to look at putative interactions between autosomal dominant and recessive alleles of genes implicated in the PD model. Comparing *park^25^/Lrrk^ex1^* with *park^25^*/+, *Lrrk^x1^*/+ and PINK1^RV^, we found that *park^25^/Lrrk^ex1^* flies show significantly reduced mobility within a minute of introduction into the arena, replicating aspects of the clinically observable bradykinesia phenotype in PD patients. Since this locomotor defect in *park^25^/Lrrk^ex1^* flies could in fact be due to the transheterozygote condition of two genes, *parkin* and *Lrrk*, implicated in PD, a detailed investigation of this possible genetic interaction will be important. Taken together, the high resolution analysis of climbing behaviour presented here helps in dissecting the possible locomotor defects of these fly mutants.

We also characterised a fly proprioceptive mutant, *Trp-γ^1^* for gravitaxis impairment. *Trp-γ*, a TRPC channel (26), is known to be expressed in thoracic bristles and femoral chordotonal organs and *Trp-γ^1^* mutants show only fine motor control defects (3). Interestingly, these flies manifested numerous descending tracks along with the ascending tracks and this was not because of less overall movement of these flies. This prompted us to investigate the geotactic ability of these mutants. To do so, we introduced the geotactic index as a way to measure fly’s response to gravity during locomotion. Graviception in flies plays an important role in vertical climbing of flies, and the geotactic index provides a way to measure and assess the innate geotactic response of the fly. Wild type *Drosophila* are sensitive to gravity and show negative geotactic response to gravity while climbing. Negative geotaxis in *Drosophila* is mediated by *TRPA* genes *pyrexia* and *painless* (27), but a role of a TRPC channel, *Trp-γ*, in gravity sensing was unknown. We found that the geotactic index of *Trp-γ^1^* mutant flies was less negative. Data from our study, thus, indicate a putative role of *Trp-γ* in graviception in *Drosophila* and warrants further studies.

In summary, we present a novel fly assay for quantifying *Drosophila* climbing behaviour at high resolution. This assay differs from traditional climbing behaviour assays by exploiting the innate negative geotactic behaviour of flies rather than relying on physical agitation. It provides insight into various locomotor parameters which are important for a quantative characterization of fly climbing. This assay, together with the open source data analysis software, opens up possibilities of extracting parameters relevant to any specific experimental question.

## Author Contributions

A.A. designed and performed the experiments, analysed data and co-wrote the manuscript. HR. and K.V.R designed the experiments, analysed the data and co-wrote the manuscript.

## Acknowledgements

We thank the *Drosophila* community and NCBS fly facility for the generous supply of fly strains. We are grateful to Dhananjay Chaturvedi for helpful discussions and suggestions. We would like to thank mechanical and electrical workshops at NCBS for fabricating the components of the setup. A.A. was supported by Council of Scientific and Industrial Research. This work was supported by National Centre for Biological Sciences – Tata Institute of Fundamental Research.Supplementary Information available at: https://gitlab.com/amanaggarwal/fly-climb-vra

---

**Supplementary figure 1.**
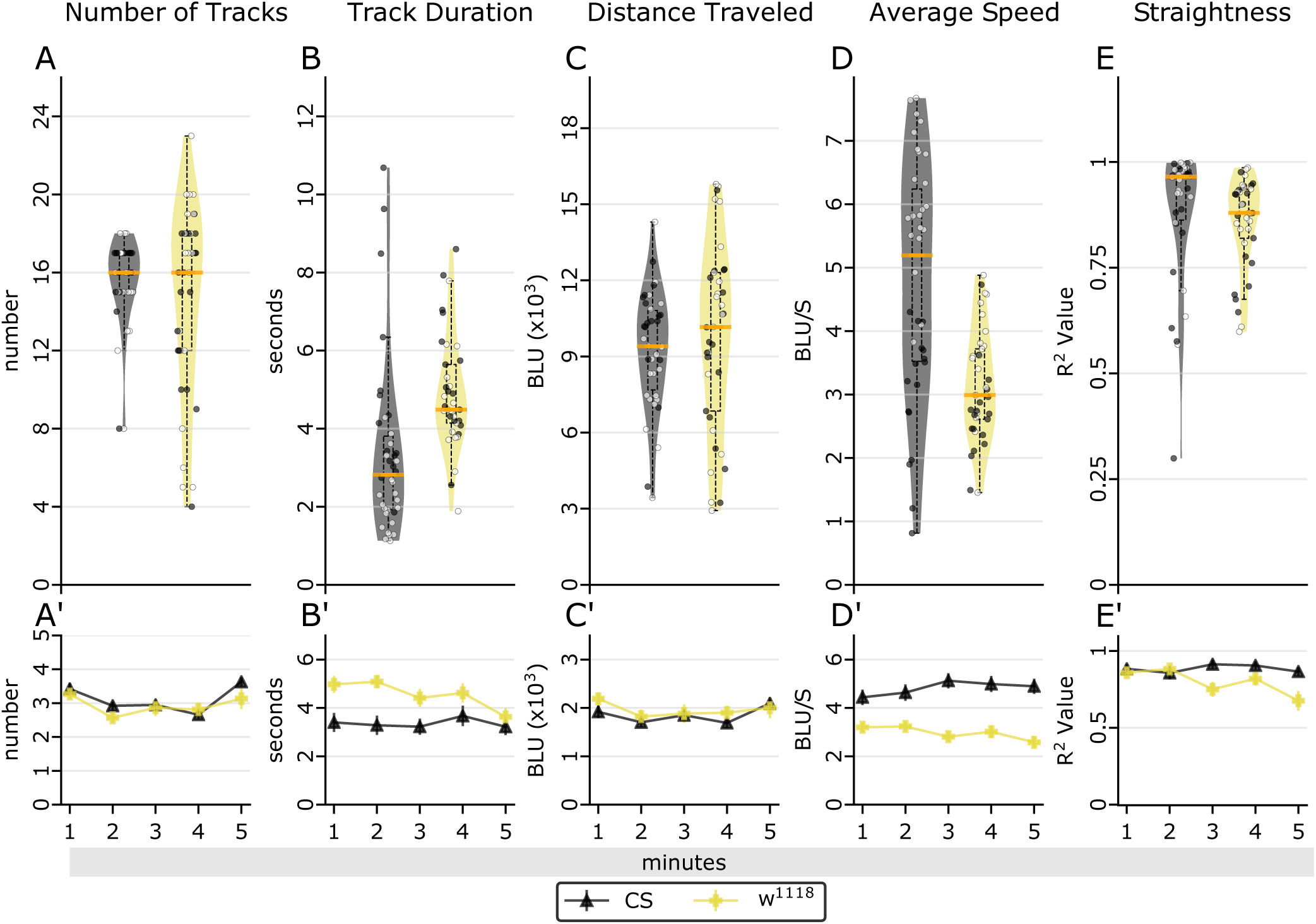
CS vs. w^1118^ flies’ climbing parameters

**Supplementary figure 2.**
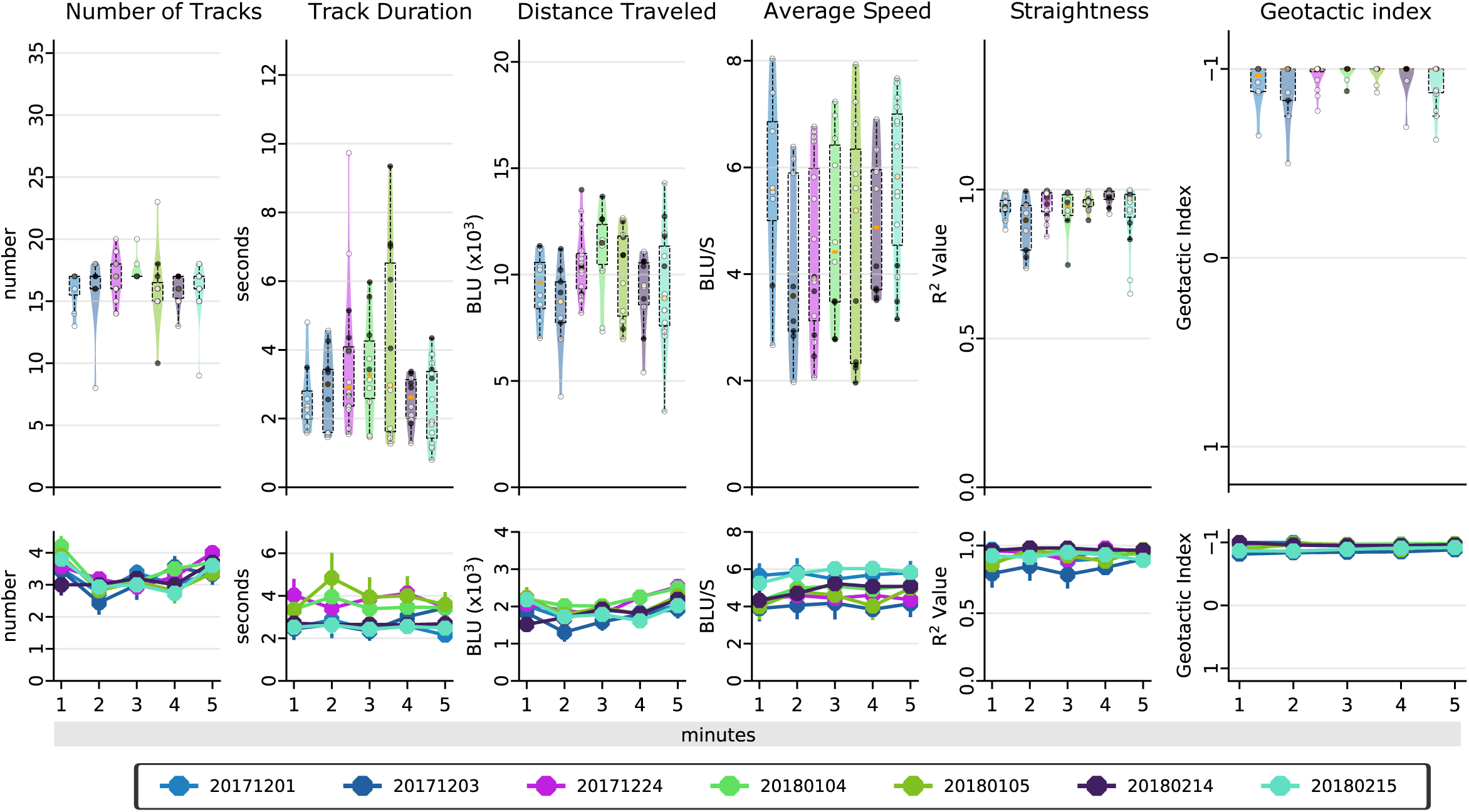
CS data compared over multiple days, showing robustness of the assay. Various climbing parameters of CS flies over different days, Each sample represents a batch of CS flies collected on a given date. date of collection is the sample name in “YYYYMMDD” format. Figure representation similar to figure 2.

